# Mouse brain-wide transgene expression by systemic injection of genetically engineered exosomes: CAP-Exosomes

**DOI:** 10.1101/2022.04.06.487362

**Authors:** Saumyendra N Sarkar, Debora Corbin, James W Simpkins

## Abstract

The bottleneck in drug discovery for central nervous system diseases is the absence of effective systemic drug delivery technology for delivering therapeutic drugs into the brain. Although some Adeno Associated Virus (AAV) serotype can cross blood brain barrier and deliver virus genome packaged therapeutic DNA (gene) or RNA molecules to brain cells along with other organs, several hurdles have emerged in the AAV9 vector gene transfer technology in both preclinical studies and clinical trials. In order to overcome some of the hurdles, we have developed a workflow to generate a novel brain targeted drug delivery system (DDS) that involves generation of genetically engineering exosomes by first selecting various functional AAV capsid specific peptides (collectively called CAP) known to be involved in brain targeted high expression gene delivery, and then expressing the CAP in frame with lysosome-associated membrane glycoprotein (Lamp2b) followed by expressing CAP-Lamp2b fusion protein on the surface of mesenchymal stem cell derived exosomes, generating CAP-exosomes. Intravenous injection of green fluorescent protein (GFP) gene loaded CAP-exosomes in mice transfer GFP gene throughout the CNS as measured by monitoring brain sections for GFP expression with confocal microscopy. GFP gene transfer efficiency is at least 20-fold greater than that of control Lamp2b-exosomes. GFP gene transduction to mouse liver was low. CAP-exosome has advantage over AAV-vector including, 1) no restriction in gene size to be delivered, 2) expected reduced production of neutralizing antibody, and 3) can be used separately for drug repurposing and/or in combination with therapeutic genes.

**Graphical Abstract:** 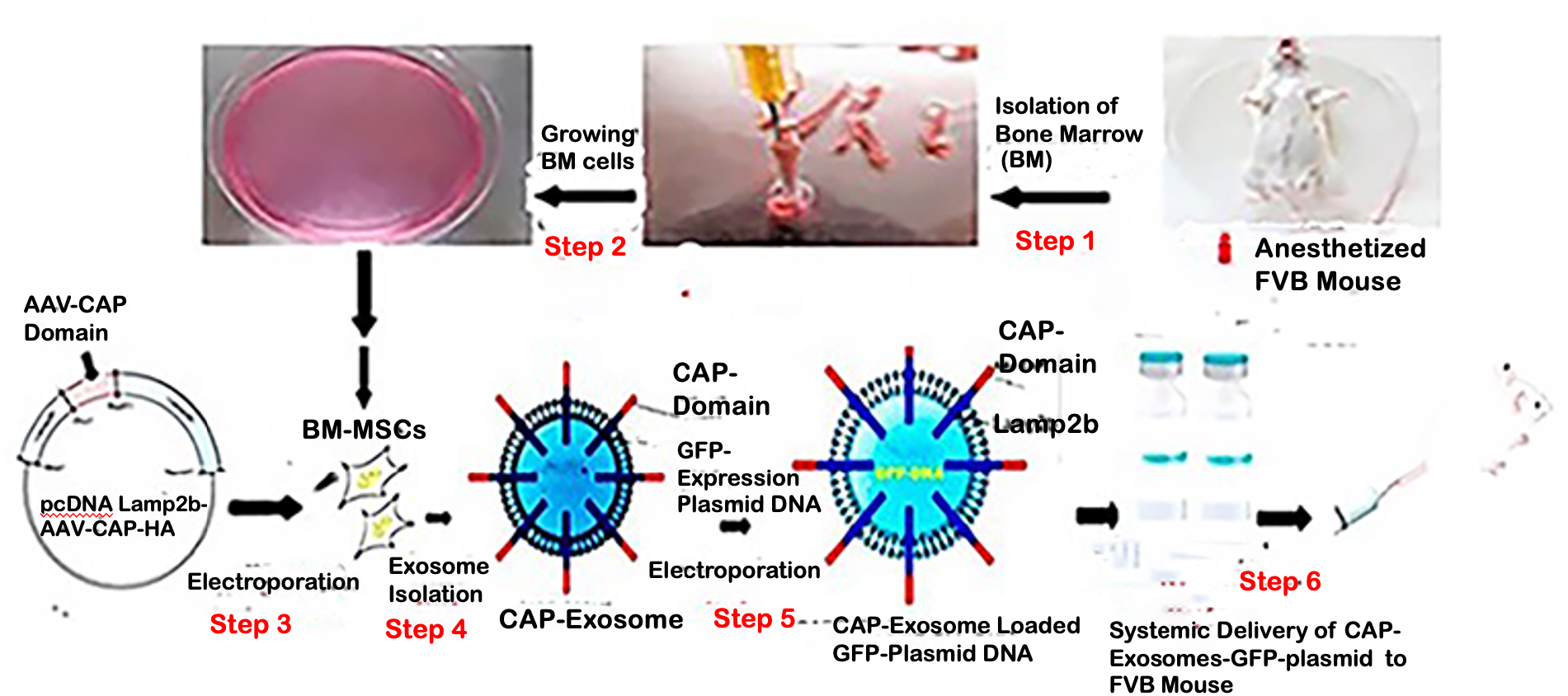

## Introduction

The aging of our population is accompanied by an increase in the absolute numbers of patients with neurological disorders, such as Alzheimer’s disease (AD) and Parkinson’s disease (PD). Specifically, Alzheimer’s disease incurs an enormous personal cost to those affected, and the worldwide financial cost in 2010 was estimated at $604 billion US ^1^.

It therefore represents a major and rising public health concern and there is an urgent need to develop more effective therapies to treat and delay the onset of the disease. Despite the advances in the technology used in drug discovery, the development of drugs for central nervous system diseases remains challenging ^2,3^. The failure rate for new drugs targeting important central nervous system diseases is high compared to most other areas of drug discovery. The main reason for the failure is the poor penetration efficacy across the blood-brain barrier (BBB). The BBB is formed by the brain capillary endothelium in the central nervous system and prevents the brain uptake by 80% of circulating small molecule drugs, and 100% of large molecule such as protein therapeutics, RNAi drugs, and other therapeutic genes.^4^

The only way the drug or gene can be distributed widely in the brain is the transvascular route following injection into the blood stream. However, this transvascular route requires the ability to undergo transport across the BBB.

Nanotechnologies such as viral and non-viral vectors allow brain targeted gene delivery systems to be created. In case of AAV nanoparticles it was discovered that certain AAV serotypes, e.g., AAV9, undergo transvascular transport across the BBB following an IV injection ^5^. AAV gene therapy of the spinal cord is now FDA approved for the treatment of infantile Spinal Muscular Atrophy (SMA) with a single IV dose of 2 × 10^14^ vector genome per kg (vg/kg) of the scAAV9 encoding the Survival Motor Neuron 1 (SMN1) gene ^6^. This gene therapeutic is designated scAAV9.CB.hSMN (Zolgensmar). The FDA approved IV dose is associated with toxicity in juvenile primates, including elevations of liver enzymes, liver failure, degeneration of dorsal root ganglia, proprioceptive deficits, and ataxia ^7^. Zolgensma IV AAV gene therapy is approved for a 1-time treatment, because subsequent doses of AAV may cause a potentially severe immune reaction due to the immunogenicity of the AAV capsid protein.^8,9^ Therefore, the current AAV vector requires improvements in transduction potency, antibody evasion and cell/tissue specificity to allow the use of lower and safer vector doses. Very recently an important capsid variant of AAV9 was generated by sequential engineering of two capsid surface exposed loops in the capsid protein, VP3. AAV particles with this variant capsid enable brain-wide transgene expression and decreased liver targeting after intravenous delivery in mouse and marmoset ^10^.

In addition to AAV vectors, exosomes, a heterogeneous group of nano-sized natural membrane vesicles, are being developed as non-viral vectors for drug delivery to the brain since they can overcome the BBB penetrance issues. However, the low targeting ability and size-dependent cellular uptake of native exosomes could profoundly affect their delivery performance. Using engineering technology, exosomes can obtain active targeting ability to accumulate in specific cell types and tissues by attaching targeting units to the membrane surface or loading them into cavities. Recent studies have indicated that brain targeted exosomes from mesenchymal stem cell (MSC) can be created by genetic engineering the exosomes with rabies virus glycoprotein (RVG) peptide expressing Lamp2b, an exosomal membrane protein. However, RVG-exosomes also transduce cells in the liver. ^11^

We reasoned that by displaying these two capsid loops in the outer membrane surface of exosomes could gain transgene-loaded exosome’s functional ability to cross BBB and allow brain-wide transgene expression without liver targeting. To this end, we engineered exosomes by first designed nucleotide sequences to encode nuclear localization signal (NLS) signal from AAV9 VP1 capsid protein, endosomal escape signal, from AAV9 VP1 capsid protein loop1, spacer, and loop2 peptides to introduce the targeting ligands at the restriction site in the N-terminus of exosomal protein Lamp2b.

We have developed a workflow to generate a novel brain targeted drug delivery system (DDS) that involves generation of genetically engineering exosomes by first selecting various functional AAV capsid specific peptides (collectively called CAP) known to be involved in brain targeted high expression gene delivery, and then expressing the CAP in frame with lysosome-associated membrane glycoprotein (Lamp2b) followed by expressing CAP-Lamp2b fusion protein on the surface of mesenchymal stem cell derived exosomes, generating CAP-exosomes.

Intravenous injection of a green fluorescent protein (GFP) gene loaded CAP-exosomes in mice transferred GFP gene throughout the CNS as measured by monitoring brain sections for GFP expression with confocal microscopy. GFP gene transfer efficiency is at least 20-fold greater than that of control Lamp2b-exosomes. GFP gene transduction to mouse liver was low.

## Results

### Characterization of Genetically Engineered CAP-Lamp2b-Hemagglutinin (HA)-Exosomes and Lamp2b-Exosomes

We have developed a workflow to generate a novel brain targeted drug delivery system (DDS) that involves generation of genetically engineering exosomes by first selecting various functional AAV capsid specific peptides (collectively called CAP) known to be involved in brain targeted high expression gene delivery, and then expressing the CAP in frame with lysosome-associated membrane glycoprotein (Lamp2b) followed by expressing CAP-Lamp2b fusion protein on the surface of mesenchymal stem cell derived exosomes, generating CAP-exosomes. (Fig 1A). For this study, a recombinant plasmid expression vector was constructed by designing nucleotide sequences to encode targeting ligand CAP domain which consists of nuclear localization signal (NLS) from AAV9 capsid protein VP1^12^, endosomal escaping signal from VP1.^13^ heptamer amino acid insertion sequences in variable region VIII of AAV9 ^14^, and spacer amino acids, heptamer amino acid substitution in variable region IV of AAVCAPB22 ^10^. CAP domain specific forward and reverse stranded DNA oligos were synthesized and obtained from Integraded DNA Technology (IDT, USA). After annealing the oligos, double stranded CAP domain was ligated at the restriction site BsaB1 in frame with the N-terminus of exosomal protein Lamp2b in pcDNA-GNSTM-3-FLAG10-Lamp2b-HA—plasmid (Hemagglutinin, HA). The pcDNA-GNSTM-3-FLAG10-CAP-Lamp2b-HA generated plasmid (Supp. Fig 1) in this study and control plasmid pcDNAHygro-Lamp2b (Addgene#86029) were used for electroporation of bone marrow-derived mesenchymal stem cell (BM-MSCs) to produce CAP-Lamp2b fusion protein and Lamp2b positive MSC-derived genetically engineered exosomes. To validate whether the CAP-lamp2b-HA and the Hygro-Lamp2b plasmids were successfully electroporated into BM-MSCs, and in case of pcDNA-GNSTM-3-FLAG10-CAP-Lamp2b-HA plasmid, the expression of CAP-Lamp2b-HA fusion protein, we assessed the protein level of Lamp2b and CAP-Lamp2b-HA fusion protein in BM-MSCs 48 hours after electroporation. Compared with BM-MSCs electroporated by pcDNA-Hygro-Lamp2bplasmid, the western blotting showed that the AAV-CAP-Lamp2b-HA transfected cells expressed HA-Lamp2b fusion protein (Fig. 1B, compare Lane 1 in both right and left diagram) if the CAP domain was not cloned at the N-terminal in frame with Lamp2b, then HA expression will be out of frame and as a result both Lamp2b and HA antibody specific signal could not be detected in the same protein by western blotting. Next, we made large-scale tissue culture growth of BM-MSCs, and electroporation with the CAP-Lamp2b-HA and Hygro-Lamp2b plasmid DNA and from culture supernatant of BM-MSCs respective exosomes were purified by ultracentrifugation. Western blot analysis of a total of 10^9^ exosomal particles (average size of 100-150 nm) showed that both Lamp2b and HA specific signal are present in Fig. 1C (Lane 1 indicating the CAP-Lamp2b-HA fusion protein inclusion on the exosomes).

**Figure 1.**
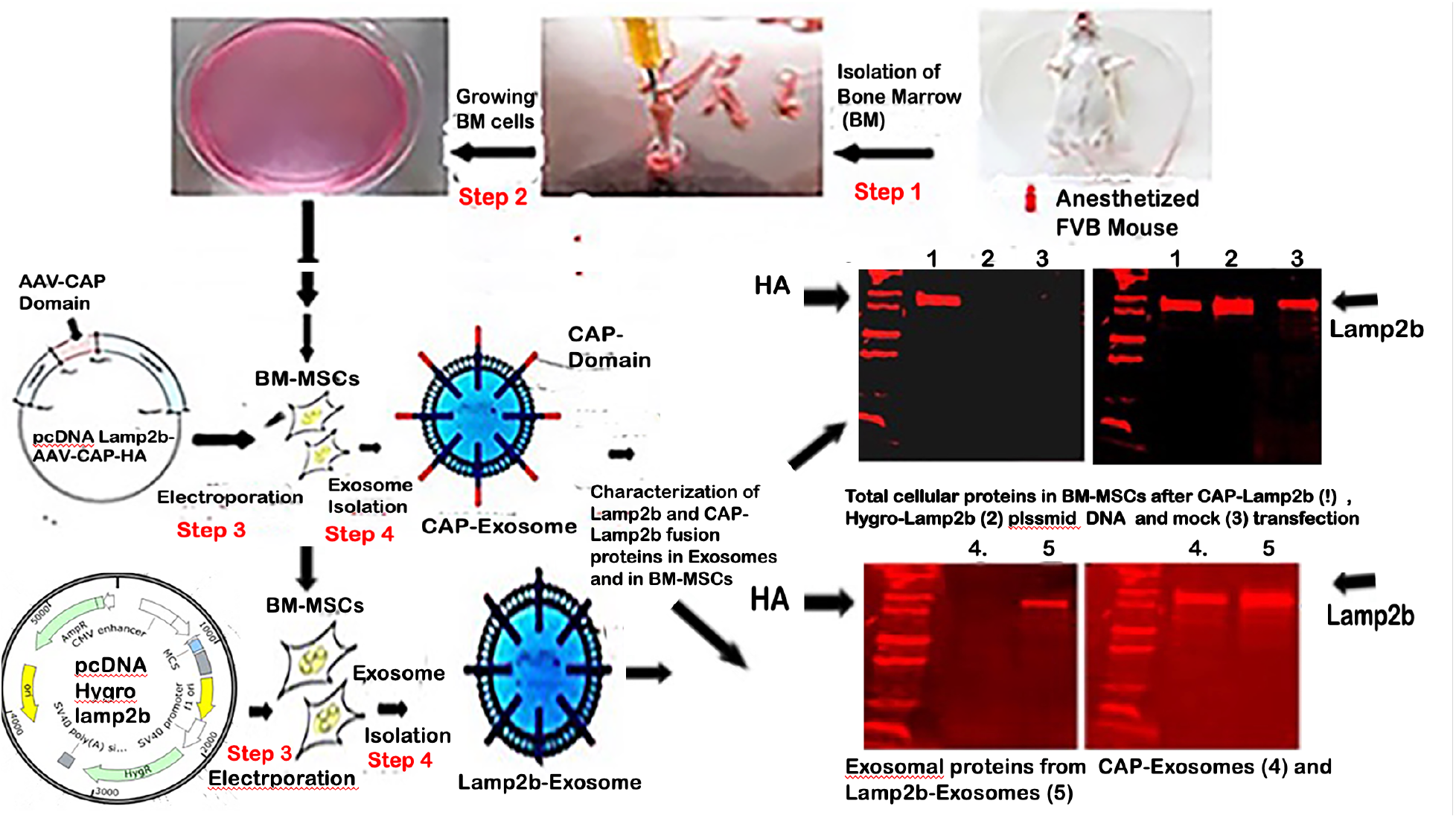
Generation and characterization of genetically engineered MSCs-derived exosomes. A). The schematic diagram showing the various steps including isolation and growth of BM-MSCs derived-AAV-CAP domain-Lamp2b fusion protein expression plasmid recombination, generation of BM-MSC derived CAP-Exosomes and Lamp2b-exosomes. B). Western blot analysis shows expression of fusion protein CAP-Lamp2b-HA in total protein isolated 36 hours after electroporation of CAP-Lamp2b with BM-MSCs as both the Hemagglutinin HA-tag antibody specific and Lamp2b-specific positive signal shows in the 1^st^ western blot run (Fig.1B. Right panel in lane1) and after striping of 2^nd^ western blot (Fig.1B, Left panel, lane1) respectively, and neither in the protein isolated from pcDNA-hygro-Lamp2b electrporated BM-MSCs (Fig.1B, Lane 2, Right and Left panel), nor in the total protein isolated from just BM-MSCs (Fig1B, Lane 3, Right and Left panel). C). Western blot analysis of a total of 10^9^ exosomal particles (average size of 100-150 nm) showed that both Lamp2b and HA specific signal are present in Fig. 1C (Lane 1) indicating the CAP-Lamp2b-HA fusion protein inclusion on the exosomes.

### Targeting efficiency and specificity of CAP-Exosomes and Lamp2b-exosomes in the brain

To investigate the potential of CAP-exosomes to specifically deliver a transgene. (GFP-gene expression plasmid), we have developed a workflow shown in Fig. 2A. First CAP-Exosomes and Lamp2b-Exosomes were labeled with 1,1’-dioctadecyl-3,3,3’,3’-tetramethylindocarbocyanine perchlorate (DiI) dye. To wash away free DiI, exosomes were resuspended in 1x phosphate-buffered saline (1XPBS) and pelleted by ultracentrifugation. Next, pelleted DiI labelled exosomes were resuspended in electroporation buffer and for a single IV injection into a FVB mice a total of 50μg of exosomes corresponding to ∼ 5×10^9^ particles were mixed with 50μg of GFP-expression plasmid DNA in a final volume of 100 μL electroporation buffer. GFP expression plasmid was inserted into respective exosomes using 4-D-Nucleofector (instruments and reagents from Lonza, Basel, Switzerland) following the company’s instructions. Mice were sacrificed 5 days after IV injection and serial coronal sections were made and images were taken by confocal microscopy.

**Figure 2.**
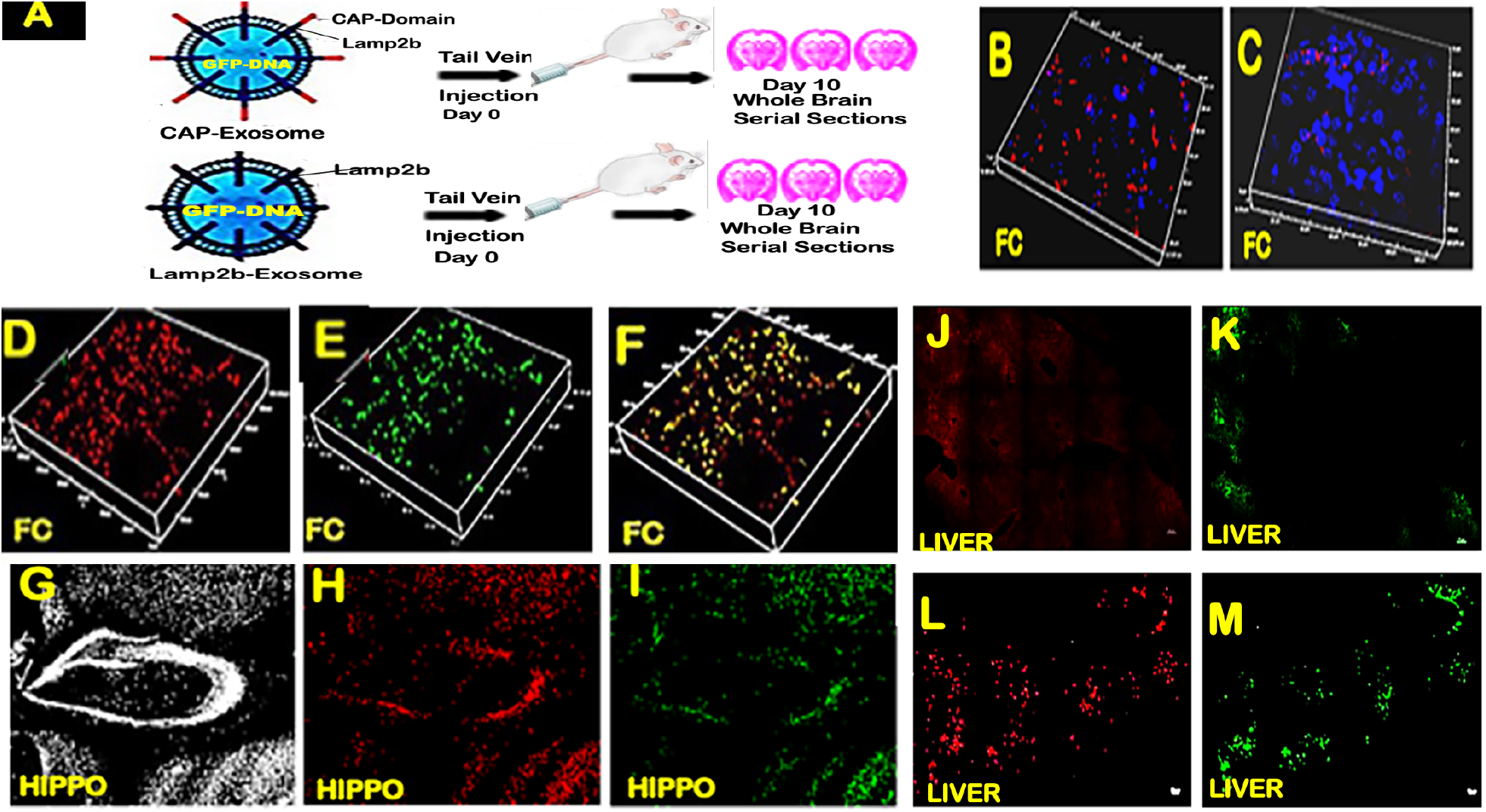
Targeting efficiency and specificity of CAP-Exosomes and Lamp2b-exosomes after systemic delivery. Presence of high no of DiI-labeled (red) CAP-exosomes (Fig. 2B) compared to Lamp2b-exosmes (Fig. 2C) n the frontal cortex region of respective mice. In the frontal cortex (FC) of the mice very high DiI-labeled exosomes were found (3D-confocal micrograph Fig. 2D, red fluorescence) along with high efficiency GFP-transgene expression (Fig. 2E, green fluorescence). Note that visual inspection of merging (Red+Green) confocal micrograph (Fig. 2F) will reveal that high percentage of only red fluorescence but not merged. Comparative studies of NeuN stained micrograph (Fig. 2G, white) with DiI labeled exosomes (Fig. 2H, red) and GFP-expression (Fig. 2I, green) indicate that systemically delivered CAP-Exosomes loaded GFP-transgene capable of brain wide and highly neuron specific delivery of genes and its expression in the mouse brain. Both the red (Fig. 2J) and green (Fig. 2K) fluorescence were low indicating CAP-domain expressing exosomes minimizing off-target GFP-gene expression. In a separate sets of animals IV injection of DiI labeled and GFP loaded hygro-Lamp2b exosomes to mice resulted in high exosomal uptake (Fig. 2L) and GFP-gene expression (Fig. 2M) in the animals liver.

In order to detect the presence of the exosomes derived from hygro-Lamp2b plasmid transfected MSCs in the brain and whether CAP-Exosomes modification enhanced the BBB crossing and enter into the brain cells, the slides of brain sections were observed under fluorescence microscope at 5 days after injection. The DiI-labeled exosomes were found in the frontal cortex of both MSC-Lamp2b-Exo group (Fig. 2C) and MSC-CAP-HA-Exo group. But when further compared by the number of DAPI-stained nuclei and DiI stained exosomes red fluorescence, there was much more DiI-labeled exosomes in the FC of the mice in the MSC-CAP HA-Exo (Fig. 2B) group than that in the MSC-Lamp2b-Exo group (Fig. 2C).

### Systemic delivery of CAP-Exosomes loaded GFP-gene resulted in brain-wide GFP expression and decreased liver targeting

In order to determine the CNS and liver targeting efficiency of CAP-exosomes various brain regions and liver sections were made from the same mice sacrificing 10 days after IV injection of DiI labeled and GFP loaded exosomes and analyzed by confocal microscopy. In the frontal cortex (FC) of the mice, very high DiI-labeled exosomes were found (3D-confocal micrograph Fig. 2D, red fluorescence) along with high efficiency GFP-transgene expression (Fig. 2E, green fluorescence). Further analysis by visual inspection of merging (Red+Green) confocal micrograph (Fig. 2F) revealed that very high percentage of exosomes either were not loaded with GFP-plasmid during electroporation or GFP-loaded exosomes were unable to escape endosomes and failed to transfer GFP-gene to the nucleus of brain cells for transcription. In order to determine the capabilities of CAP-exosomes for neuron specific brain-wide gene transfer, brain sections from the same mice used for Fig. 2D, were stained with neuronal marker NeuN antibody and analyzed nearly the whole hippocampus regions by confocal microscopy using low power (10x) objective lens. Comparative studies of NeuN stained micrograph (Fig. 2G, white) with DiI labeled exosomes (2H, red) and GFP-expression (Fig. 2I, green) indicate that systemically delivered CAP-Exosomes loaded GFP-transgene capable of brain wide and highly neuron specific delivery of genes and its expression in the mouse brain.

For further studies to determine whether the high expression of intravenously delivered CAP-exosomes-loaded GFP gene in CNS cells while minimizing off-target expression, liver sections obtained from the same mice as was used in for Fig. 2 and were analyzed by confocal microscopy. Both the red (Fig. 2J) and green (2K) fluorescence were low indicating CAP-domain expressing exosomes minimizing off-target GFP gene expression. In a separate sets of animals IV injection of DiI labeled and GFP loaded hygro-Lamp2b exosomes to mice resulted in high exosomal uptake (Fig. 2L) and GFP-gene expression (Fig. 2M) in the animals liver.

To further characterize brain specific GFP-gene delivery by CAP-exosomes compared with Lamp2b-exosomes, we stained for neurons with NeuN and DAPI nuclei stained for liver cells and quantified the gene transfer efficiency and target specificity of each exosome for each cell type across various brain regions and in the liver (Fig. 3. A and B). Quantification of the total number of neuron specific NeuN stained ells expressing GFP in the Hippocampus shows 9% of neurons were expressing GFP in case of CAP-exsomes injected mice verses 1.5% in case of control exosomes. In the cortical region of mice injected with CAP-exosomes 20% of neurons were GFP positive compared with 2% in case of control exosomes. Whereas neurons were expressed GFP gene with high efficiency in CAP-exosomes injected mice compared with control exosomes, GFP expression in liver cells 5-6-fold lower levels in CAP-exosomes treated mice compared with control exosomes. This indicates that the CAP-exosomes have neuronal specificity in mice.

**Figure 3.**
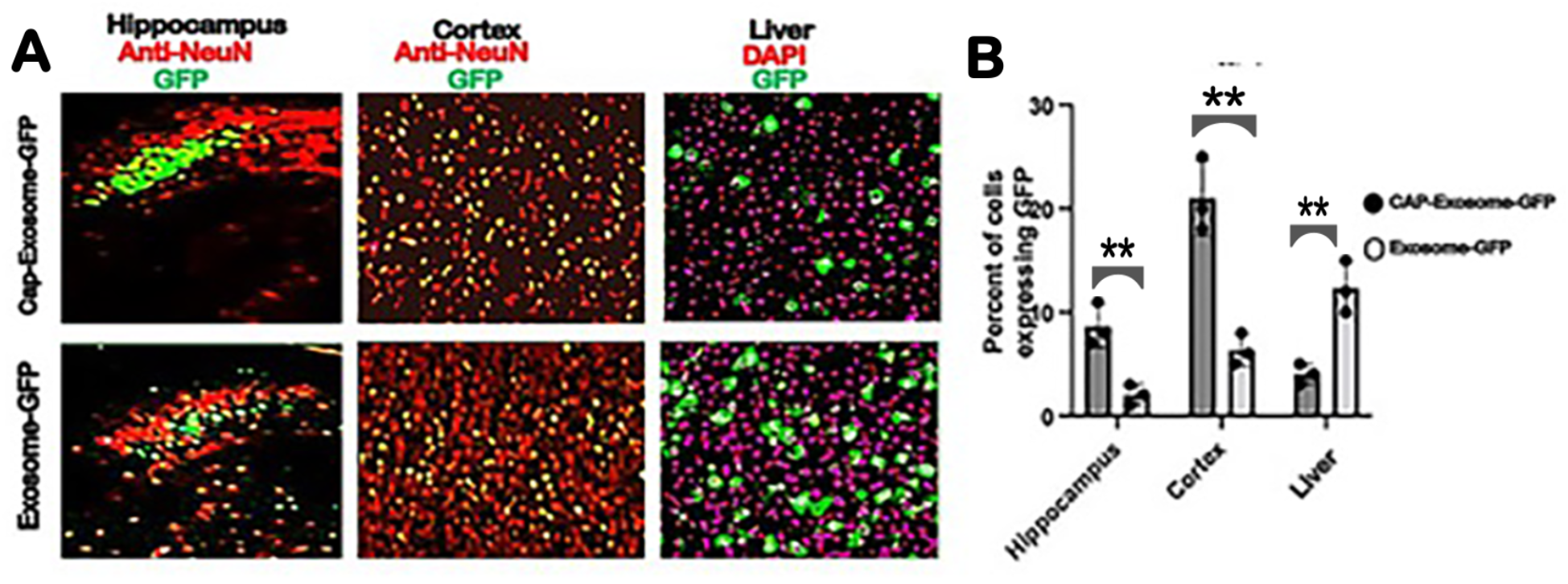
CAP-Exosome-GFP expression is preferred in brain with a significant decrease in liver. CAP-Exosome-GFP (5μg in o.1 ml saline buffer/mice) and Exosome-GFP (5μg in o.1 ml saline buffer/mice) particles were intravenously injected into FVB adult mice. GFP fluorescence was assessed after one week of expression. Expression of the total number of cells expressing GFP and NeuN immune stained shown in (A) and percentage of Cells expressing GFP shown in (B) in the Hippocampus (n=4 animals *P* = 0.0074, control Exosome versus CAP-Exosomes), Cortex (n=4 animals, p = 0.002, control Exosome versus CAP-Exosomes). In case of Liver total number of cells expressing GFP and DAPI nuclear stained cells shown in (A) and percentage of cells expressing GFP shown in (B) (n=4 animals, p= 0.005, control Exosome versus CAP-Exosomes).

## Discussion

The goal of this study was to use genetic engineering of exosomes to confer its ability to cross the BBB efficiently and then further refine exosomes for delivery of drugs to neurons. To achieve this special properties by exosomes, we first chose two AAV9 variant capsid surface exposed loops specific peptide along with NLS and endosomal escaping signal peptide as ligand (we called this peptide as CAP-domain) and fused with transmembrane proteins Lamp2b that are expressed on the surface of the exosomes. Subsequently, BM-MSCs were chosen as donor cells and transfected with plasmids encoding the CAP-Lamp2b fusion proteins engineered exosomes, CAP-exosomes bearing targeting ligands on their surface membrane. We chose CAP-domain as the ligand because it has been reported that these two variant surfaces exposed peptide loop structure is responsible for conferring AAV’s ability to cross BBB highly efficiently and this variant capsid enable brain-wide transgene expression and decreased liver targeting after intravenous delivery in mouse and marmoset.^10^

Intravenous injection of green fluorescent protein (GFP) gene loaded CAP-exosomes in mice transfer GFP gene throughout the CNS as measured by monitoring brain sections (hippocampus and frontal cortex) for GFP expression with confocal microscopy. GFP gene transfer efficiency was at least 20-fold greater than that of control Lamp2b-exosomes. GFP gene transduction to mouse liver was low. Also, we chose expression vector that is capable of producing glycosylation-stabilized CAP-domain-Lamp2b fusion protein and is important for increasing the expression of CAP-domain on exosome ^15^. The strategy used in this study for generating CAP-exosomes may be particularly useful for translating promising exosome-based drug delivery from preclinical investigation to human trials for brain diseases. Currently the only one phase I/II clinical trial to explore the safety and efficacy of Adipose-MSCs-exosomes in the treatment of mild to moderate dementia due to Alzheimer’s disease. (NCT04388982)

Mesenchymal stem/stromal cell-exosomes are an emerging treatment for a variety of inflammatory and degenerative conditions and are beginning to be translated from preclinical models into early phase clinical trials ^16^. Engineering exosomes for targeted drug delivery increases the local concentration of therapeutics and minimizes side effects ^17^. Although still in its infancy, as the CAP-exosome-mediated drug delivery boasts low toxicity, low immunogenicity, and capabilities of brain wide delivery of transgene, CAP-exosomes holds promise for nano-particle mediated therapies for a wide range of CNS diseases.

## 2. Materials and methods

### Animals and cell culture

All rodent procedures were approved by the Institutional Animal Use and Care Committee of West Virginia University.

### FVB mice bone marrow cell collection

The anesthetized FVB mice (2-3 months old) was placed in a 100-mm culture dish, and washed with 70% ethanol. Tibias and femurs were dissected; muscle, ligaments, and tendons were removed. Next, with micro dissecting scissors, the two ends were excised and a needle with the syringe filled with sterile phosphate buffered saline was inserted into the bone cavity and used to slowly flush the marrow out into a culture dish containing BM-MSC growth medium. BM-MSC colony growth was expanded according to the protocol described by Masoud Soleimani & Samad Nadri ^18^ and used for transfection of CAP-Lamp2b, and hygro-Lamp2b expression cassette containing plasmid DNA by electroporation. GFP expression plasmid was inserted into CAP-exosomes using 4-D-Nucleofector (instruments and reagents from Lonza, Basel, Switzerland, following the company’s instructions).

### Construction of CAP-Lamp2b expression vector

Nucleotide sequences were designed to encode targeting ligand named CAP domain which consists of <u>NLS signal</u>_from AAV9 capsid protein VP1, endosomal escaping signal from VP1, heptamer amino acid insertion sequences in variable region VIII of AAV9, spacer<u>_</u>amino acids, heptamer amino acid substitution in variable region IV of AAVCAPB22. CAP domain specific forward and reverse stranded DNA oligos were synthesized and obtained from Integraded DNA Technology (IDT, USA). After annealing the oligos, double stranded CAP domain was ligated at the restriction site BsaB1 in frame with the N-terminus of exosomal protein Lamp2b in pcDNA-GNSTM-3-FLAG10-Lamp2b-HA—plasmid (addgene#71293). The pcDNA-GNSTM-3-FLAG10-CAP-Lamp2b-HA generated plasmid in this study and control plasmid pcDNAHygro-Lamp2b (addgene#86029) were used for electroporation of bone marrow-derived mesenchymal stem cell (BM-MSCs) to produce CAP-Lamp2b fusion protein and Lamp2b positive MSC-derived genetically engineered exosomes. Nucleotide sequences encoding variant AAV9-capsid surface exposed peptide loops used in this study is shown below:

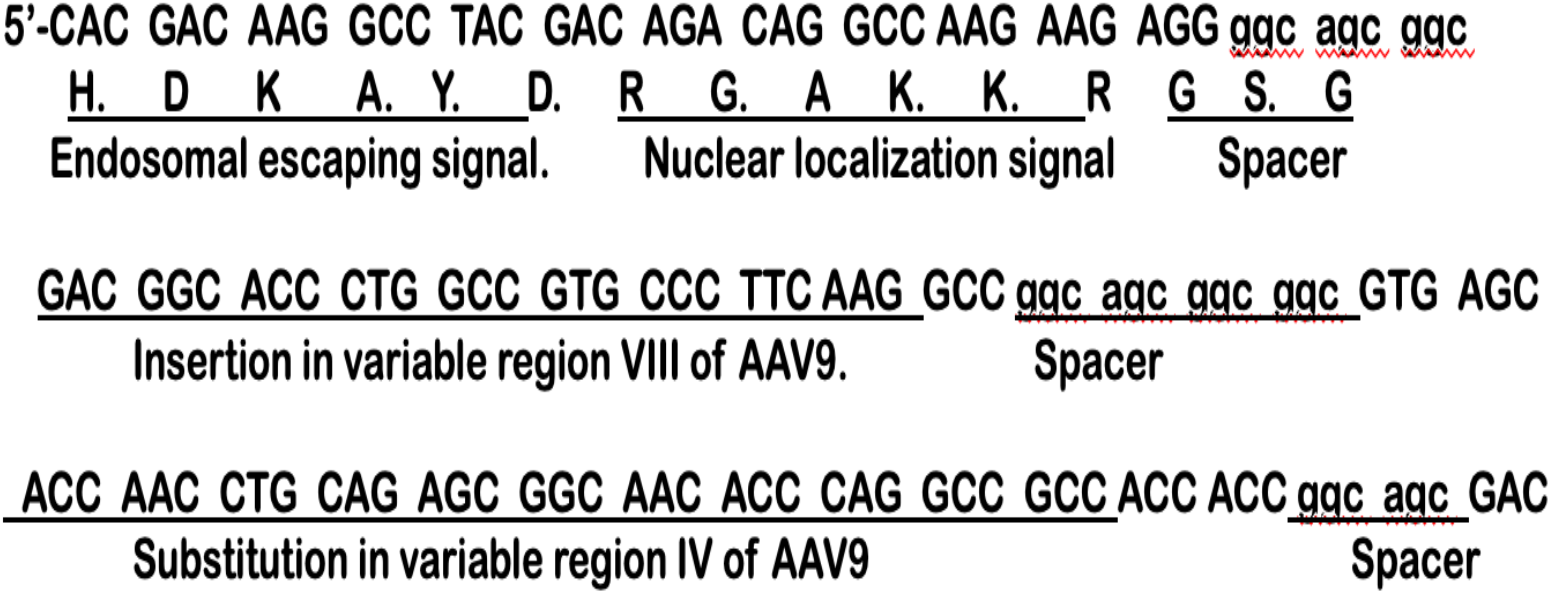

### Isolation of Exosomes

From CAP-Lamp2b or hygro-Lamp2b plasmid DNA transfected BM-MSC cell culture conditioned media exosomes were isolated and purified either by differential centrifugation as described in by Thery et al. ^19^ or by using the ExoEasy Maxi Kit (Qiagen)

### Size and concentration determination of exosomes

Sizes and concentration of purified and DiI labelled exosomes were determined by using Malvin Panalytical NanoSight instrument (USA) according to the company’s instructions.

### Western Blotting

For cell western blotting, BM-MSCs electroporated with pcDNA-hygro-Lamp2b or CAP-Lamp2b-HA-plasmid were lysed by RIPA buffer. Blots were incubated with primary antibodies overnight at 4C. The primary antibodies used were as follows: goat anti-Lamp2b (1:1,000, Abcam), and rabbit anti HA antibody (Cell Signaling Technology). Corresponding IR Dye secondary Donkey anti goat and Donkey ant rabbit (1:20,000, Lico were incubated for 1 hr at room temperature. Bands were visualized using Licor imaging system.

### Immunofluorescence

Serial coronal sections of 40 micron in thickness were prepared on a cryostat. After being incubated with 0.3% Triton X-100 and 3% bovine serum albumin (BSA) in PBS for 1 hr, the following primary antibody was incubated overnight at room temperature: rabbit anti-NeuN (1:200, Abcam 177487). The corresponding secondary antibodies was Alexa Fluor 647 (1:200, Thermo Fisher Scientific, A32795) was incubated for 3 hr at room temperature. Cellular nuclei were stained by Hoechst 33342 (1:100, Sigma).

### Imaging and Quantification

All GFP green expressing tissues (Liver and Brain), presence of DiI (red) labelled exosomes in the tissues and NeuN immunostained neurons in brain sections were imaged on a NICON A1R confocal laser Microscope using 10x and 40x objective, with matched laser powers, gains and gamma across all samples of the same tissue section. In all cases in which fluorescence intensity was compared between samples, exposure settings and changes to gamma or contrast were maintained across images. The acquired images were processed using NIS-Element imaging software.

All CAP-Exosome-GFP and Lamp2b-GFP expressing tissues were imaged on a Zeiss LSM confocal microscope using a 10x objective, with matched laser powers, gains, and gamma across all samples of the same tissue. The acquired images were processed in Zen Bue 2 (Zeiss). Processing of all czi images from confocal microscope was performed with FIJI (ImageJ). Colocalization between the GFP signal and NeuN antibody or DAPI staining was performed using FIJI with the cell count automated plugin. The details method used for the quantitation was followed according to the published method ^20^.

### Statistics

GraphPad Prism 9 were used for statistical analysis and data representation. All experimental groups were n = 4. For the statistical analyses and related graphs, a single data point was defined as two tissue sections per animal, with multiple technical replicates per section when possible. Technical replicates were defined as multiple fields of view per section, with the following numbers for each region or tissue of interest: cortex = 3 hippocampus = 4, liver = 4. Statistical analyses of the data was performed using Prism 9 and using the multiple unpaired (non-parametric) t-test.

## Acknowledgements

This work was supported by NIH Grants P20 GM109098, P01 AG027956, T32 AG052375 and U54 GM10494243T.

