## supplemental figure 1 for "Mouse brain-wide transgene expression by systemic injection of genetically engineered exosomes: CAP-Exosomes"

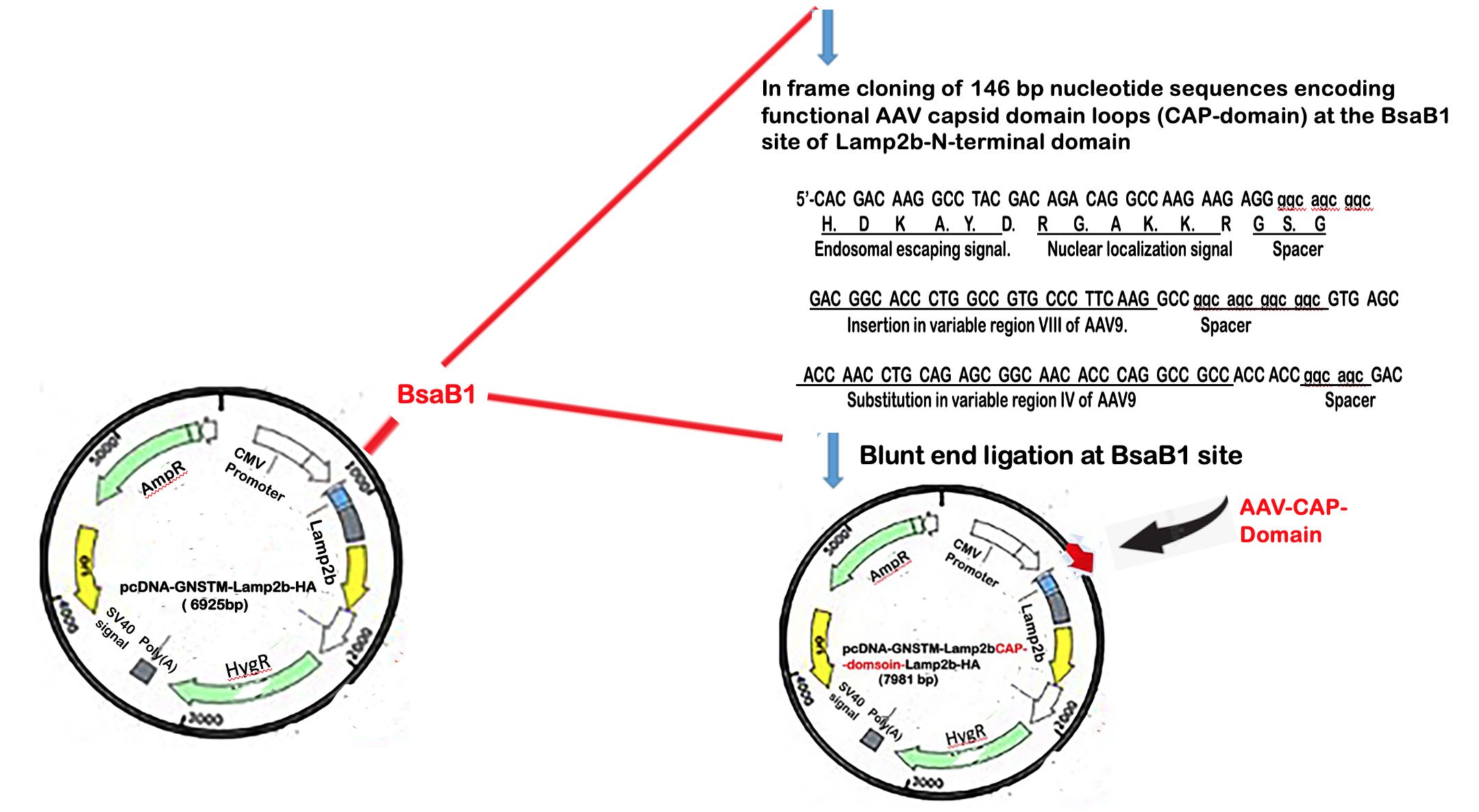

**Supplementary Fig 1:** Graphical presentation of cloning strategy for AAV-CAP domain in pcDNA GNSTM-3-Flag-10-lamp2b-HA expression plasmid vector. (Addgene, USA, Plasmid no.71293)
